# Solubility product constant directs the formation of biomolecular condensates

**DOI:** 10.1101/2020.12.26.424446

**Authors:** Aniruddha Chattaraj, Michael L. Blinov, Leslie M. Loew

## Abstract

Biomolecular condensates, formed by liquid-liquid phase separation (LLPS), are important cellular structures. Using stochastic network-free kinetic models, we establish a physical-chemical basis for the concentration threshold of heterotypic multivalent molecules required for LLPS. We associate phase separation with a bimodal partitioning of the cluster distribution into small oligomers vs. huge polymers. The simulations reveal that LLPS obeys the solubility product constant (Ksp): the product of monomer concentrations, accounting for ideal stoichiometries, does not exceed a threshold no matter how much additional monomer is added to the system – additional monomer is funneled into large clusters. The Ksp applies over a range of valencies and stoichiometries. However, consistent with the importance of disordered domains for LLPS, removing flexible linker domains funnels valency-matched monomers into a “dimer trap”, and Ksp no longer defines a threshold for large cluster formation. We propose Ksp as a new tool for elucidating biomolecular condensate biophysics.

## Introduction

Biomolecular condensates comprise a novel class of intracellular structures formed by a biophysical phenomenon called liquid liquid phase separation (LLPS) [1–3]. These structures serve as membraneless compartments where complex biochemistry can be organized and facilitated [4]; for example, T cell receptor mediated actin nucleation efficacy spikes up multifold when all the associated signaling molecules concentrate into a condensate [5] near the plasma membrane. These structures are also implicated in many age-related or neurological diseases [6–8].

Numerous theoretical and experimental studies have illuminated many of the biophysical requirements for condensate formation [9–11]. In particular, it is firmly established that clustering of weakly interacting multivalent proteins or nucleic acids is a prerequisite for the phase separation. Even a sufficiently concentrated solution of a single self-interacting protein (homotypic interaction) with multiple binding sites in its sequence, can partition into protein-dense and dilute phases. Such homotypic systems display a strict threshold concentration above which phase separation occurs. This phase separation serves as a buffering mechanism for the protein in the dilute phase [4, 12], which remains at the threshold concentration; thus, as more protein is added to the system, the dense phase droplets grow in size and number, keeping the concentration clamped in the dilute phase. Although a homotypic system closely conforms to a single fixed threshold concentration, the picture gets much more complex with multicomponent (heterotypic interactions) systems [13], which underlie all the biomolecular condensates found in living cells and contain complex mixtures of multivalent proteins and/or nucleic acid. A thorough recent study of the thermodynamics of the liquid-liquid phase transitions in heterotypic systems showed clearly that concentration thresholds for phase separation no longer remain fixed and vary with relative compositions of interacting binding partners [13]. But a theoretical framework for quantitatively predicting such complex varying concentration thresholds is still lacking.

We have previously distinguished between strong multivalent interactions, which can produce molecular machines, and weak multivalent interactions, which can produce what we termed “pleomorphic ensembles” [14, 15]. The strong binding affinities in molecular machines (e.g. ribosomes, flagella) enforce a specific parts list with a fixed stoichiometry. Pleomorphic ensembles (e.g. cytoskeletal polymers and their associated binding proteins, neuronal post synaptic densities, etc.) are much more plastic in their composition than molecular machines. Biomolecular condensates are a subclass of pleomorphic ensembles in that their molecular components simultaneously coexist within both a distinct phase and as solutes in the surrounding solution. In trying to quantitatively understand the relationships governing condensate formation from heterotypic components, we asked if there may be lessons to learn from freshman ionic solution chemistry, where salts in solution are in equilibrium with a solid crystalline phase composed of a lattice of counter-ions. We realized that the liquid phase separation at threshold concentrations of heterotypic biological molecules might resemble the precipitation of anions and cations from solution.

Consider a salt (let’s say silver chloride, AgCl) in water; dissolution takes place until the solution becomes saturated; further addition of salt results in precipitation. In the simplest case of pure AgCl, this seems to be similar to the behavior of a homotypic (single component) condensate; that is, above a threshold total concentration of dissolved AgCl, the salt will not dissolve further, maintaining a clamped concentration of Ag^+^ and Cl^−^ in the solution above, whatever amount of AgCl is added. However, the key concept is that the saturation threshold is governed by the solubility product – the product of the individual concentrations of Ag^+^ and Cl^−^, [Ag^+^] * [Cl^−^]. Precipitation starts when this product reaches the thermodynamic parameter called the solubility product constant, Ksp. For AgCl Ksp = 1.7 × 10^−10^M^2^ at 25°C. This is the basis for crystallization as a warm saturated solution cools, because Ksp usually decreases as temperature decreases. Importantly, if we add any Cl^−^ in the form of a highly soluble salt (e.g. KCl) into the solution, some AgCl will precipitate to maintain the solubility product at Ksp. Of course, the crystal lattice in an ionic solid is very different from the set of weak multivalent interactions inside a biomolecular condensate. But we wondered whether the Ksp might at least approximately be used to understand heterotypic LLPS.

We explore how well Ksp may apply to biomolecular condensates using two stochastic modeling approaches. With a non-spatial network-free simulator (NFsim [16]), we show how Ksp can approximately predict the phase separation threshold for a two-component system by systematically changing the concentration of individual components with a variety of valencies. We then test our theory with a more complex three-component system and to our gratification, the theory seems to work there as well. We then expand on prior work on the structural features governing multivalent clustering [17] with a spatial kinetic simulator (SpringSaLaD [18]); we show how molecular flexibility, valency matching, steric effects and inter binding sites distances can influence the phase separation threshold.

## Materials and methods

### Non-spatial simulations (NFsim)

To develop models that probe for the effects of valency and concentrations but do not account for spatial effects, we employ the Network-free simulator (NFsim [16]) - a non-spatial rule based stochastic simulation framework where each biomolecule represents a molecular object that may have multiple binding sites. These sites can bind with other sites depending on a set of rules defined in the model. The simulations have units of molecular counts, but these can be readily converted to equivalent concentrations, which is how we present our results.

The NFsim model file is specified in BioNetGen Language, or BNGL (http://bionetgen.org/). Let’s take the example of tetravalent binders – A_4_ (a1, a2, a3, a4) and B_4_ (b1, b2, b3, b4). We need 16 binding rules to define all the bimolecular interactions, each having an affinity (Kd) of 350 μM. Now 1000 molecules of A_4_ and B_4_ with an affinity of 3500 molecules would translate to 100 μM molecular concentrations with 350 μM binding affinity. There are two equivalent ways to change the molecular concentrations – 1. Change the Kd, keeping the molecular counts same, which is mathematically equivalent to changing the volume of the system; 2. Change the molecular counts, keeping the Kd same. We utilize both approaches for our simulations and specify which is used in our descriptions of the results. The binding rules only allow inter-molecular binding; internal bond formation within the molecular clusters is not permitted, as NFsim cannot account for proximity of binding sites within clusters. Once the BNGL file is defined, we then generate the corresponding XML file which serves as the NFsim input file. We run multiple stochastic simulations in parallel using the high performance computing facility at UConn Health (https://health.uconn.edu/high-performance-computing/). A single NFsim run (500 ms, fixed total concentration approach), containing 1000 A_4_ and B_4_ molecules each, took ~ 1 min with 100 trajectories run in parallel. A python script is then used to perform statistical analysis across all the trajectories.

### Spatial simulations (SpringSaLaD)

To account for realistic spatial geometry, we employ SringSaLaD [18] - a particle based spatial simulation platform where each biomolecule is modelled as a collection of hard spheres connected by stiff spring-like linkers. The simulation algorithms are fully described [18] and we previously studied various spatial biophysical factors in the context of multivalent biomolecular cluster formation [17] with this software. SpringSaLaD also uses a rule based method to define binding reactions between multivalent binders.

The SpringSaLaD model files are generated using the graphical user interface (GUI) of the software (https://vcell.org/ssalad). We define the size of binding sites, distance between the binding sites and the overall shape of the molecule inside the GUI. If we take the example of reference system (A_4a_ and B_4b_), two linear tetravalent molecules are constructed first, each having four binding sites and six linker sites. Unlike NFsim, in SpringSaLaD, one binding rule between “A_type” and “B_type” sites can take care of all the possible binding interactions. Also we can define the binding affinity in concentration units (350 μM) directly inside the GUI. We initialize our system in a 3D rectangular geometry; for example, 100 molecules in a 10^6^ nm^3^ cubic volume (X = Y = Z = 100 nm) would correspond to 166 μM. We change the volume of our system to alter the molecular concentrations. Once the model is specified, as before, we run multiple stochastic simulations in parallel using our high performance computing facility. Execution time is very much sensitive to the number of total sites due to the computational overhead of tracking individual site locations. A typical run (50 ms, fixed total concentration approach), containing 100 molecules each of A_4a_ and B_4b_ (total sites = 2000), took ~ 6 hours.

### Data Analysis and Visualization

Python scripts are used to analyze and visualize the data. All the scripts are written with Spyder IDE (version 4.0.0) https://www.spyder-ide.org/. Frequently used python libraries are numpy 1.17.3, pandas 0.25.3, matplotlib 3.1.2. All the packages are managed by anaconda package distributions https://www.anaconda.com/.

All the model files, Python scripts and a “Readme” description of all the contents are available in a public GitHub repository: https://github.com/achattaraj/Ksp_phase_separation.

## Results

### Phase boundary of a two-component system with molecules of the same valency

Our baseline model system consists of a pair of tetravalent molecules (A_4_ and B_4_), each having four binding sites that can interact with an affinity (Kd) of 350 μM. We start the NFsim simulation with the same concentrations of monomers of each type and measure the free monomer concentrations when the system reaches the steady state (Fig. 1A). The product of both free concentrations is called solubility product (SP). As we increase the total concentration (synchronously of both A_4_ and B_4_), the concentration of monomeric molecules initially goes up (Fig. 1A insert) and of course SP also goes up; upon reaching a concentration threshold (total concentration ~60 μM in this case), SP converges to a constant value (169 μM^2^). This is the solubility product constant or Ksp for this pair of tetravalent molecules.

**Figure 1:**
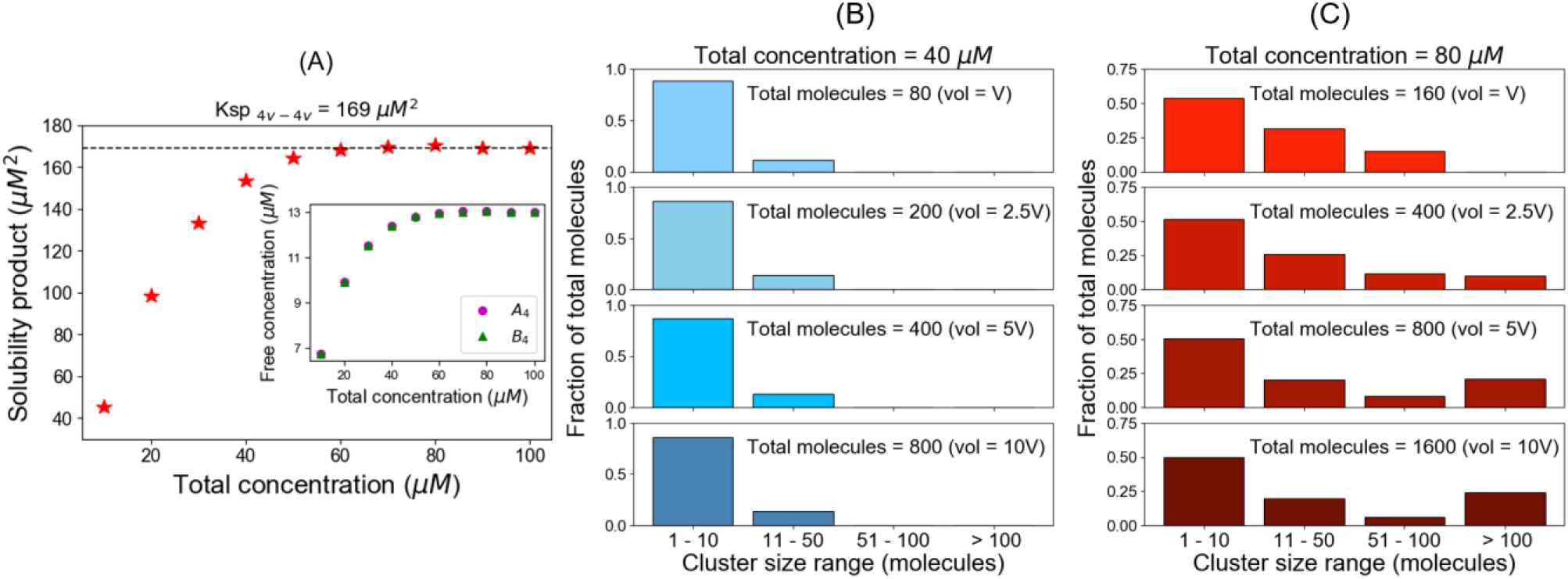
The solubility product constant corresponds to a threshold above which molecules distribute into large clusters. These simulation results correspond to equal total concentrations of a heterotypic tetravalent pair of molecules with Kd for individual binding of 350 μM. (A) Product of the free monomer concentrations as a function of the total molecular concentrations. The black dashed line indicates the plateau, corresponding to the solubility product constant (Ksp), 169 μM^2^. Inset plot shows the change of free molecular concentrations of both tetravalent molecules with their respective total concentrations. Each data point is an average of steady state values from 200 trajectories. In these simulations, we titrate up the molecular counts (100, 200, 300, …., 1000 molecules respectively), keeping the system’s volume fixed. (B, C) Distribution of cluster sizes with varying system sizes at two different total concentrations, 40 μM and 80 μM, respectively below and above the plateau in (A). The histograms show how the molecules are distributed across different ranges of cluster sizes.

Next, we ask how the system might behave differently below and above the Ksp. To test that, we analyze the cluster size distributions at two different total concentrations, 40 μM and 80 μM (Figs. 1B, 1C). At each of these two concentrations, we probed for how the number of available molecules (i.e. the system volume) affects the cluster size distribution.

Fig. 1B illustrates the cluster distribution for the 40 μM total concentration (SP = 154 μM^2^ < Ksp). The shape of the cluster size distribution displays an exponential decline from monomers to higher oligomers and this shape is insensitive to increasing the number of molecules (i.e. volume) in the system (unbinned histograms are in Supp. Fig. S1). Even in the presence of 800 molecules, there are hardly any clusters greater than 40 molecules (lowest panel of Fig. 1B). Almost 80% of the total molecules reside in the smallest cluster bin, which is 1-10 molecules per cluster. Extrapolating to a macroscopic system, this would be equivalent to a single soluble phase consisting of mainly monomers and small oligomers.

Fig. 1C illustrates the cluster distribution for total concentration = 80 μM (SP = 169μM^2^ = Ksp). The histogram does change shape, becoming bimodal as we feed more molecules into the system (i.e. as we increase the volume). With 1600 molecules in the system, more than 20% of the total molecules are in clusters larger than 100 molecules. Supp. Figs. S2 (panels A and B reproduce the same behavior with a much larger system (10,000 total molecules). After 60 μM molecular concentrations, SP converges to a plateau (Ksp) and cluster size distribution more prominently becomes bimodal. Thus, Figs. 1C and S2B demonstrate in two different ways that if the system is above Ksp, molecules are funneled into macro-clusters.

We hypothesize that this tendency to form increasingly larger clusters with more available molecules is a hallmark of phase separation behavior and that a constant solubility product (Ksp) is a quantitative indicator to mark the threshold for phase separation that underlies biomolecular condensates. Below a threshold total concentration, when SP hasn’t reached the constant level of Ksp, the tendency to form large clusters is low and the system would exist as a single phase (e.g. Fig. 1B). Above the threshold where SP converges to the Ksp, the system tends to form very large clusters, yielding two different phases, manifest as bimodal cluster size histograms (Fig. 1C and Supp. Fig. S2B). The dense phase containing the larger clusters grows in size and the dilute phase concentration remains constant. Importantly, this behavior is precisely that of a buffering system, which has been proposed as one of the important biophysical functions of biomolecular condensates. The generality of this hypothesis will now be further explored through additional modeling scenarios.

### Simulations with monomeric A_4_ and B_4_ maintained at fixed concentrations

We now demonstrate that Ksp marks the phase transition threshold using an alternative modeling approach. In Fig. 1, we used a fixed total concentration (FTC) of molecules and measure the free monomer concentrations as the system reaches the steady state. In this second approach, we clamp the monomers concentration to a constant value (“CMC”, Fig. 2A) and allow the total concentration (free plus bound) to change over time. This is achieved by creating reactions that rapidly create and destroy monomers, such that the concentration is clamped at the ratio of these rate constants - as long as these rates are much faster than the rates of the binding reactions. The solubility product, in this case, is simply the product of CMC_A_4_ and CMC_B_4_.

**Figure 2:**
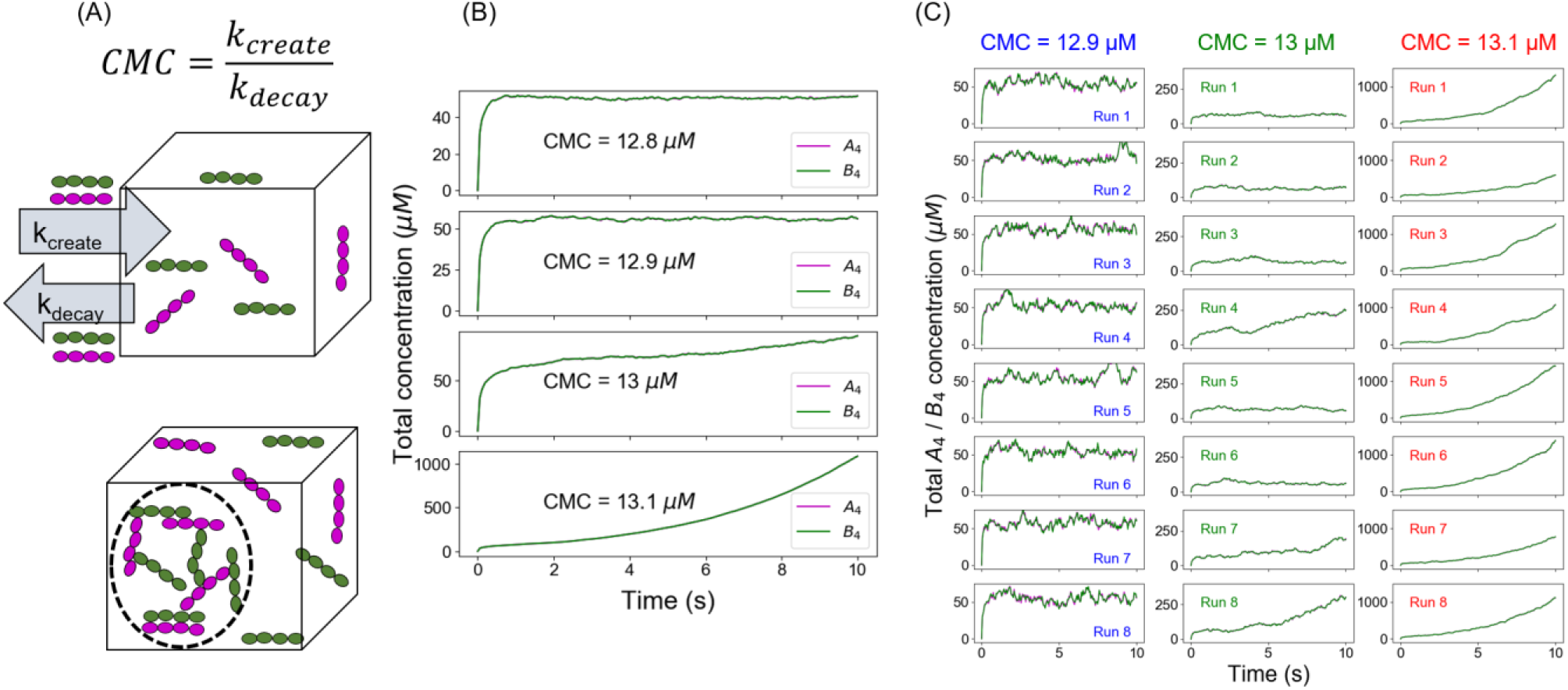
An alternative approach to quantify the phase transition boundary. (A) Illustration of the clamped monomer concentration (CMC) approach. Both the molecules (A_4_ in magenta and B_4_ in green) can enter the simulation box with a rate constant k_create_ (molecules/s) and exit with a rate constant k_decay_ (s^−1^). The ratio of these two parameters clamps the monomer concentration to (k_create/_ k_decay_). (B) Average time course (over 100 trajectories) of total molecular concentrations as a function of different CMCs. (C) Eight sample trajectories for different CMCs.

For CMC_A_4_ = CMC_B_4_ = 12.8 μM (SP = 163.8 μM^2^, below Ksp), total molecular concentrations rise up initially and then converge to a steady state (~ 50.6 μM) (Fig. 2B). Similar behavior can be observed for CMC = 12.9 μM where the total concentration converges to 56 μM. However, going to CMC = 13 μM (SP = 169 μM^2^ = Ksp), the total concentrations never reach a steady state. This phenomenon is more pronounced for a higher CMC (13.1 μM). Clearly the system is undergoing a fundamental change around SP = 169 μM^2^, identical to the threshold determined for FTC (Fig. 1). Both these modeling paradigms indicate that there is a solubility product constant (Ksp = 169 μM^2^) beyond which the system has a much higher propensity to form larger molecular clusters, the prerequisite for phase separated droplet formation. Each panel in Fig. 2B is the average of 100 trajectories; when we look at the individual trajectories (Fig. 2C and Supp. Fig. S3), the system shows much larger stochastic fluctuations near the phase boundary (Fig 2C, CMC = 13 μM); the total concentration explodes in some runs, or remains steady around a stable state within the given timeframe. The behavior is less stochastic away from the Ksp (Fig 2C, CMC = 12.9 μM and 13.1 μM). The behavior at Ksp represents stochastic nucleation of clusters containing enough crosslinking that disassembly becomes unlikely; such larger clusters are only probable at or above Ksp.

When we compare the outcomes from FTC and CMC methods, we see that the results are consistent with each other (Figs. 1A, 2B and Supp. Fig. S4). Below the phase boundary, if we clamp the monomer concentrations to the value of free concentrations obtained from the FTC method, we recover the same steady state total molecular concentrations (Top 6 panels in Supp. Fig. S4). However, at even slightly above the Ksp, the CMC total concentration increases with time, rather than leveling off to a higher steady value. (Bottom panels in Supp. Fig. S4).

### Phase transition depends on Ksp even when individual monomer concentrations are unequal

From ionic solution chemistry, we know that irrespective of the individual ionic concentrations, if the product of ion concentrations exceeds the Ksp of the salt (i.e. supersaturation), we always get precipitation to restore the solution concentrations to Ksp, even if one ion is present at a different concentration than the other (“common ion effect”). We wanted to test whether that simple chemical principle works for our relatively complex multivalent molecular clustering system. In Fig. 3, we analyze two cases when product of reactant’s concentrations (solubility products, SP) is above (SP = 172 μM^2^) and below (160 μM^2^) the Ksp (169 μM^2^, derived from Fig. 1). For each of those SPs, we vary the CMCs of A_4_ and B_4_ in such a way that the products of the CMCs are always equal to the assigned SP. Satisfyingly, we find that for SP < Ksp (Fig. 3B), the systems always converge to stable steady states (no phase transition); but for SP > Ksp (Fig. 3A), irrespective of individual clamped concentrations, systems always show unbounded growth (phase transition). We also titrated unequal fixed total concentrations, maintained at a ratio of 3:2 (supp. Fig. S2, panels C and D). Interestingly, in this computational experiment the free monomer concentration of the lower abundant component (B_4_) gets exhausted disproportionately as the threshold is approached, so that Ksp cannot quite reach 169 μM^2^ and the SP actually begins to diminish somewhat at still higher FTC. However, cluster size distribution (Supp. Fig. S2D) still becomes increasingly bimodal with higher concentrations suggesting a phase separating behavior. Thus, even when monomer supply becomes depleted, the Ksp still serves as an upper limit for free monomer concentrations.

**Figure 3:**
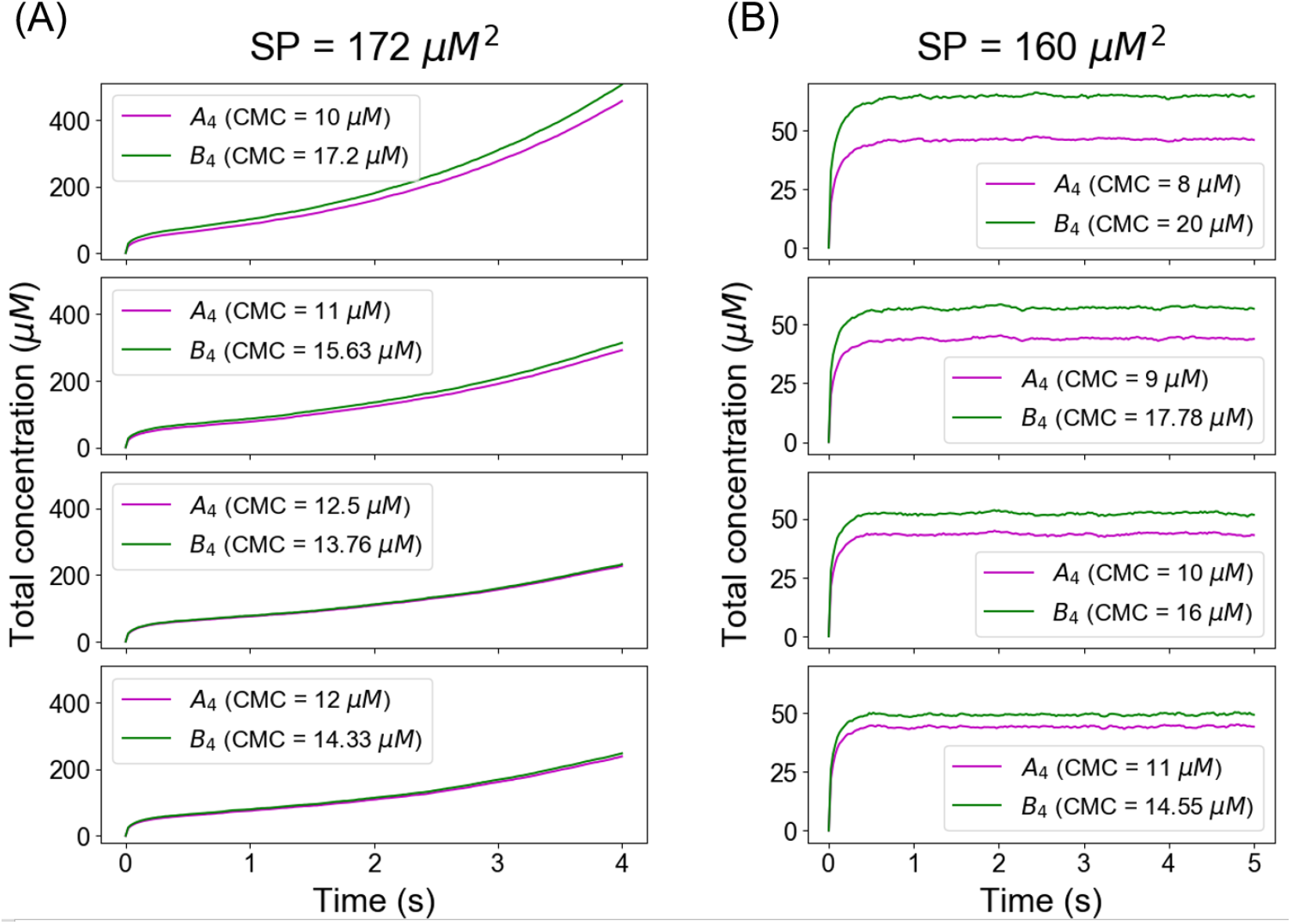
The Ksp defines a threshold for unlimited growth of clusters even when the individual concentrations of heterotypic multivalent binding partners are unequal. With Ksp determined from Figs 1 and 2 at ~169 μM^2^, SP was clamped above in (A) at 172 μM^2^ and below in (B) at 160 μM^2^.

### Higher valency promotes phase transition by reducing the Ksp

Increasing valency is known to increase the propensity for phase separation [4, 8]. Therefore, we ask how the valency of the interacting heterotypic monomers affects the Ksp. We altered the molecular valencies (number of binding sites per molecule) from 3 to 5 and compute the solubility product profiles as a function of total concentrations (Fig. 4). The total concentration needed to reach the Ksp goes down with higher valency, generally consistent with experiment. Going from 3v,3v pair to 4v,4v pair, Ksp changes over 5 fold (852 μM^2^ to169 μM^2^), whereas ~ 3 fold change (169 μM^2^ to 55 μM^2^) can be observed for transitioning into 5v,5v pair from 4v,4v.

**Figure 4:**
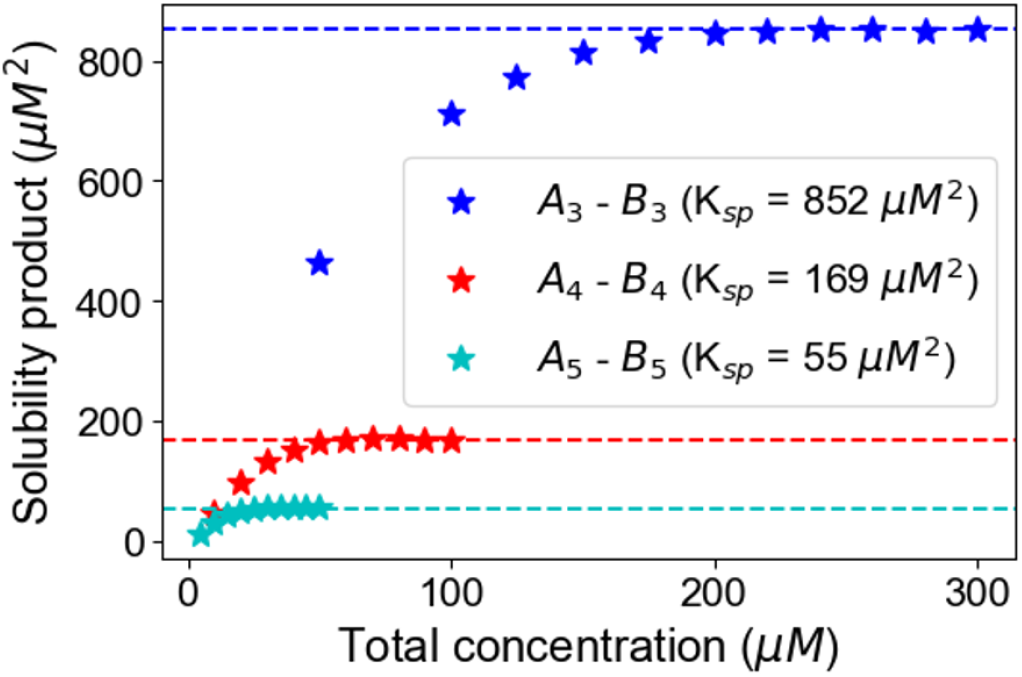
Change of Ksp with molecular valency. For 3,3 case (blue stars), molecular counts of each type = [500, 1000, 1500, …, 3000]. For 4,4 case (red stars), molecular counts of each type = [100, 200, 300, …, 1000]. For 5,5 case (cyan stars), molecular counts of each type = [50, 100, 150, …, 500]. Kd is set to 3500 molecules in all these cases. Horizontal dashed lines indicate the Ksp of the corresponding system.

### Ksp for a mixed-valent system

We next explore what happens when we mix molecules with different valencies. Consider a penta- and tri-valent (A_5_-B_3_) molecular pair (Fig. 5). To optimize the clustering such that all sites could potentially be bound would require a stoichiometry of 3A_5_ : 5B_3_. Maintaining this concentration ratio, as we titrate up the total concentrations, we see an interesting pattern (Fig. 5A, inset): the free monomer concentration of B_3_, which is present in excess, goes up steadily; but the free A_5_ goes up first and then starts to go down. When we take the product of free monomer concentrations (Supp. Fig. S5), we don’t see a solubility product profile that plateaus to a constant Ksp (as in Fig. 1A). However, when we correct the solubility product expression by taking the ideal stoichiometry into account, SP = (free A_5_)^3^(free B_3_)^5^, we get a SP profile that does plateau to a fixed Ksp beyond the total concentration threshold of ~128 μM (Fig. 5A). The Ksp expression for this mixed-valent binary system is analogous to a mixed-valent salt like Al_2_ (SO_4_)_3_. Indeed, examining the cluster size distribution confirms that this mixed-valent system has a concentration threshold around 128 μM (48 μM A_5_ + 80 μM B_3_); beyond that point, SP converges to a constant value, the cluster size distribution becomes bimodal and more and more molecules populate the larger clusters (Fig. 5B). Thus, the Ksp analogy between ionic solution chemistry and biomolecular condensates seems to hold for even these more complex stoichiometries.

**Figure 5:**
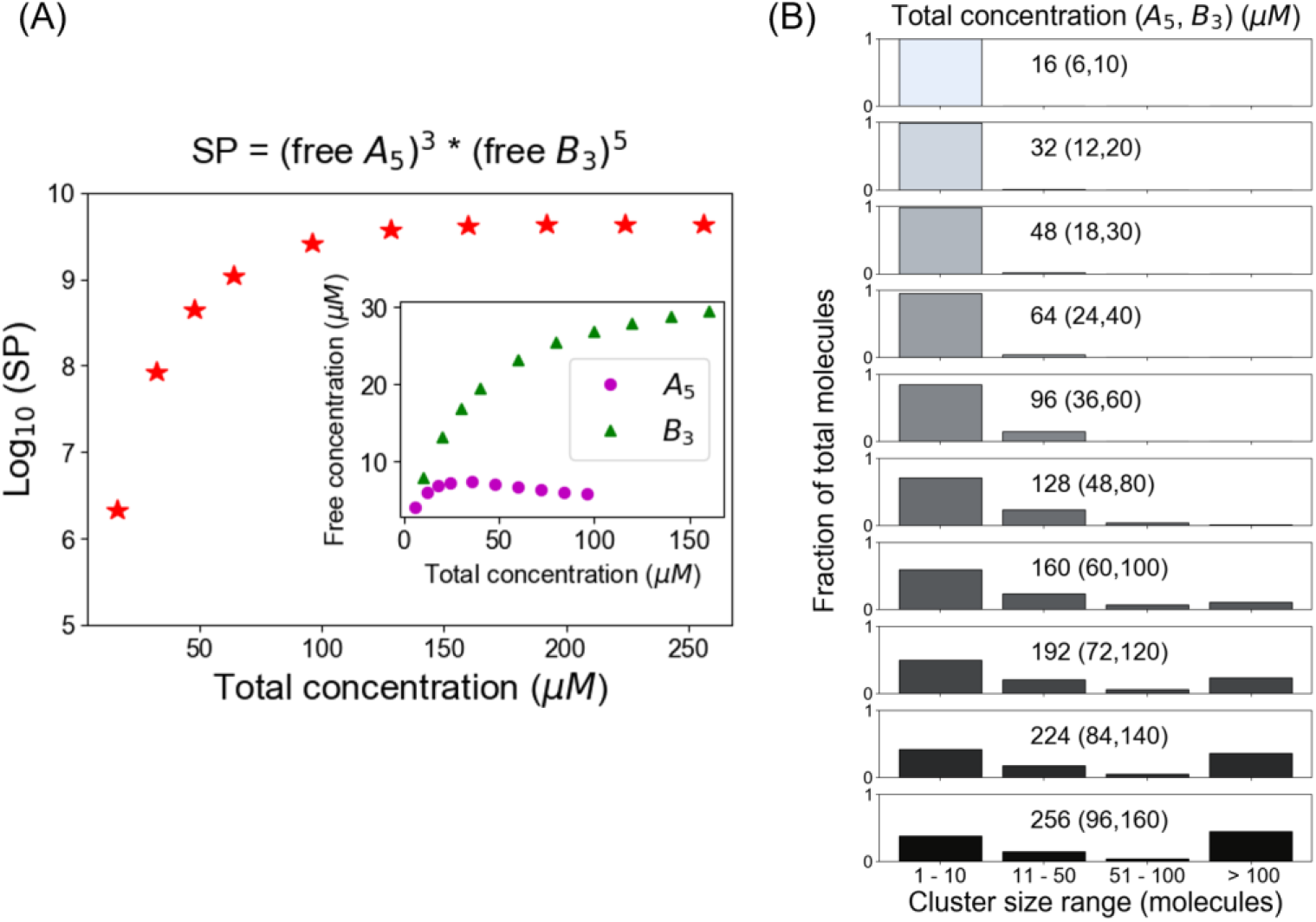
Mixed valent binary system follows the same Ksp expression as of a mixed-valent salt like Al_2_ (SO_4_)_3_. (A) Logarithm of SP as a function of total concentrations (A_5_ + B_3_). Inset shows the variation of free molecular concentrations w.r.t their initial total concentrations. Molecules are added at fixed Kd (= 3500 molecules) to change the concentrations (B) Cluster size distributions become more bi-modal as we go beyond the critical concentration (128 μM in this case)

### Ksp for a three component system

We have thus far shown that the idea of a solubility product constant or Ksp seems to work well for a variety of scenarios involving heterotypic binary multivalent interactions. A natural question would be whether the principle can be generalized for a more complicated system. Here we consider a mixed-valent three component system, inspired by the now classical study by the Rosen lab [8] on the Nephrin-Nck-NWASP (Fig. 6A) binding partners. Nephrin has three phospho-tyrosines which can bind to a single SH2 domain of Nck; six proline-rich domains (PRMs) in NWASP interact with the three SH3 domains of Nck. In our simulations, all these bindings are assumed to have the same weak affinities (Kd = 350μM). The initial molecular concentrations are chosen to maintain matching numbers of binding sites of each type; for example, 2 molecules of Nephrin with 6 phosphotyrosine sites require 6 Nck molecules, each with one SH2 site; 6 molecules of Nck (each with 3 SH3 sites) would require 18 PRMs or 3 NWASP molecules (each with 6 PRMs). So the molecular stoichiometry is 2 Nephrin : 6 Nck : 3 NWASP for perfect binding. Keeping that ratio of initial molecular counts, we vary the concentrations and quantify the free monomer concentrations at steady state (Fig. 6B, inset). We compute the solubility product (SP) as a function of these three free molecular concentrations, raised to their respective stoichiometries. It is satisfying to see that the SP profile of this complex three-component system reaches a threshold, as in the binary systems, corresponding to a Ksp in this case of ~10^12^ μM^11^. This plateau is reached at a total concentration (sum of the three total concentrations) of 165 μM (Fig. 6B); above this threshold, the histogram separates into a bimodal distribution (6^th^ panel of Fig. 6C), indicative of phase separation. Thus, even for this more complex mixed valent ternary system, a threshold for monomer concentrations is set by a Ksp, and it determines the point for formation of the large cluster phase.

**Figure 6:**
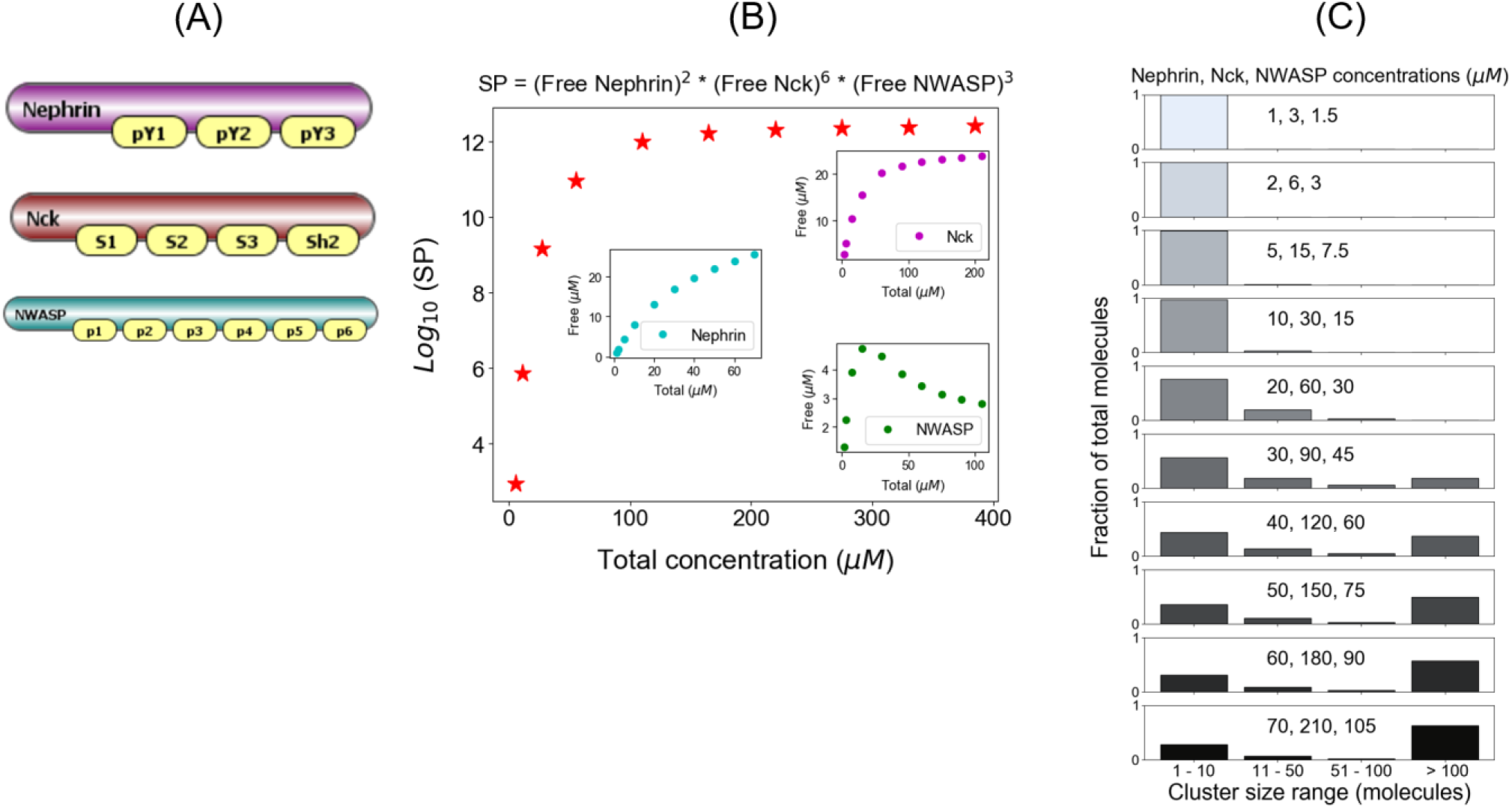
Ksp for a three component system. (A) Nephrin is a transmembrane protein which has three tyrosine residues (pY1, pY2, pY3) at its cytoplasmic tail; when phosphorylated, it can bind to the SH2 domain of a signaling protein called Nck. Nck also has three SH3 domains (S1, S2, S3 in the model) which can recruit another signaling protein called NWASP through the binding of six proline rich motifs (p1, p2, p3, p4, p5, p6). Binding affinities for both interaction types are low; Kd _(pY-Sh2)_ = Kd_(S-P)_ = 350μM. (B) SP is plotted on log scale as a function of total molecular concentrations. Inset shows the variation of free concentrations of individual molecules. Molecular counts of Nephrin, Nck and NWASP were set to 200, 600, 300 molecules respectively; we change the Kd to alter the molecular concentrations (C) Cluster size distributions at various molecular concentrations. The distribution starts to become more bimodal around 165 μM (30, 90, 45 μMs).

### Spatial simulations

Because of the efficiency of the NFSim non-spatial stochastic simulator, we were able to rapidly explore many scenarios using large numbers of molecules and demonstrate that the solubility product constant (Ksp) may generally serve as a quantitative indicator for phase transitions of multivalent heterotypic binders. These simulations also allowed us to focus on only the effects of valency and stoichiometry. We now apply a spatial simulation framework, SpringSaLaD [18], to understand the roles of spatial features, such as steric hindrance, molecular flexibility and proximity, on the Ksp. The spheres may be designated as binding sites within a rule-based modeling interface, assigning them macroscopic on and off rates that the software translates to reaction probabilities as the spheres penetrate an automatically computed reaction radius. Thus, the software is amenable to computational experiments where the geometric and reaction parameters of the system can be systematically varied.

We initially characterized a “Reference” system using the two complementary modeling approaches (FTC and CMC) to see if spatial simulations also support the applicability of Ksp. This “Reference” system consists of a pair of tetravalent molecules (A_4a_ and B_4b_) where each molecule contains four binding sites, with interspersed pairs of linker sites (Fig. 7A). These linker sites impart flexibility to the molecules mimicking the intrinsically disordered linker sequences that are found in many phase separating multivalent proteins [19]. Each of the magenta sites can bind to each of the green sites with an affinity of 350 μM. We begin with 100 A_4a_ and 100 B_4b_ molecules, randomly distributed in a three-dimensional rectangular volume. As the system relaxes to the steady state, we quantify the monomer concentrations and plot their product (solubility product) as a function of the initial total concentration – the FTC approach as we described before for NFsim. We generate a solubility product profile by systematically varying the FTC by changing the volume of the compartment, keeping the total molecular numbers same (Fig. 7B). The solubility product converges to a constant value (Ksp = 318 μM^2^) near a critical FTC (~ 138 μM in this case). When we look at the detailed cluster size distributions at steady state (Supp. Fig. S6), the dimer emerges as a preferred configuration both below and above the Ksp. This dimer preference arises from the matching valency and spatial arrangement of the binding sites in the two partner molecules; such an effect, which we term a “dimer trap” cannot be realized in non-spatial methods, such as NFsim.

**Figure 7:**
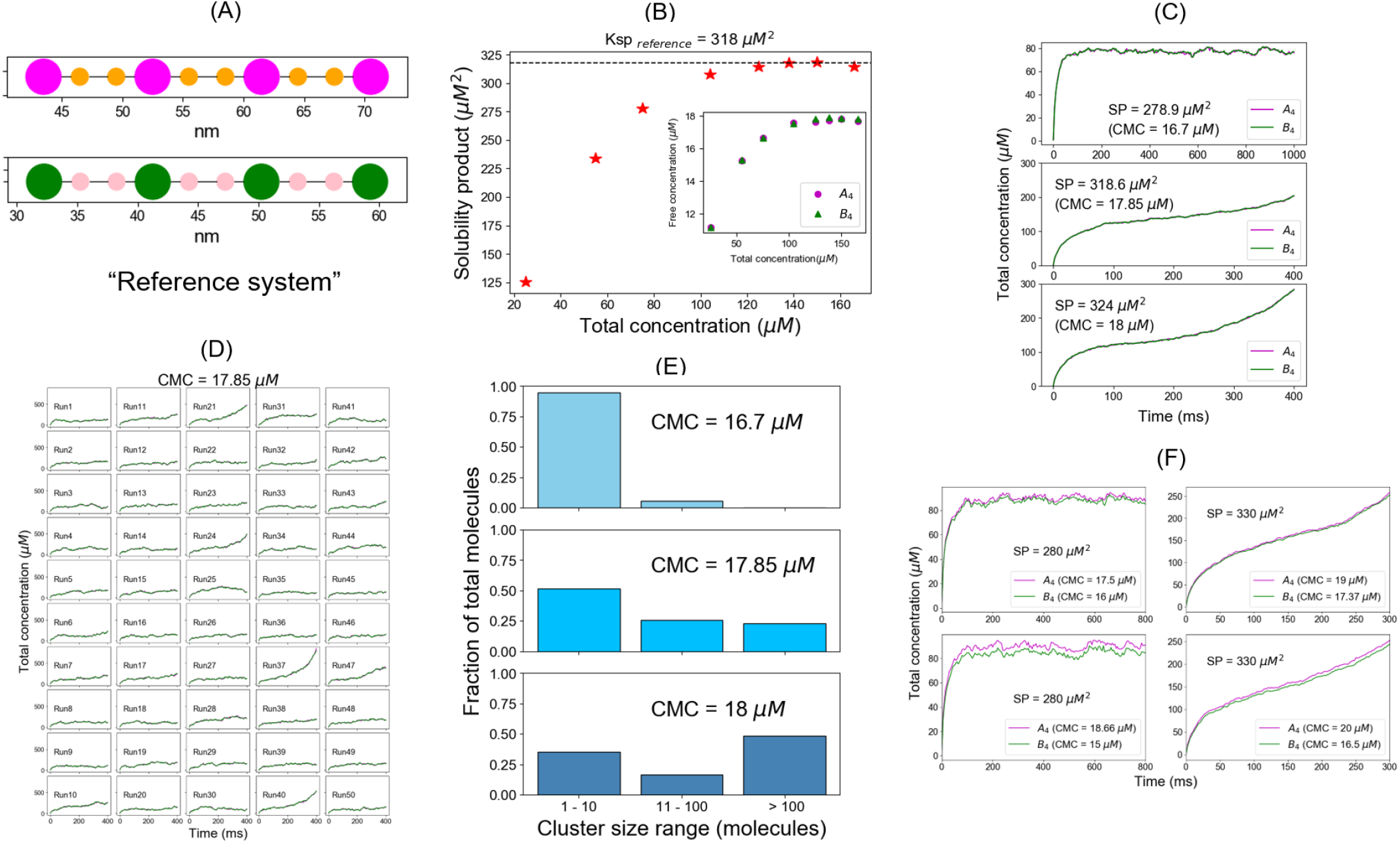
Spatial simulations demonstrate similar phase boundary behavior as NFsim. (A) SpringSaLaD representations of a pair of tetravalent binders. A_4a_ and B_4b_ consist of 4 magenta and green spherical binding sites (radius = 1.5 nm) and 6 orange and pink linker sites (radius = 0.75 nm). Diffusion constants for all the sites are set to 2 μm^2^/s. For individual binding, dissociation constant, Kd = 350 μM. We set this configuration as our reference system. (B) Solubility product (SP) profile of the reference system. We place a total of 200 molecules (100 A_4a_ + 100 B_4b_) in 3D boxes with varying volumes and quantify the monomer concentrations (free A_4a_ and free B_4b_) at steady states. Each data point is an average over 100 trajectories. Solubility product constant (Ksp) = 318 μM^2^, the horizontal dashed line. (C) Total molecular concentration profiles for three clamped monomer concentrations (CMCs). These profiles are averaged over 50 trajectories. (D) Individual trajectories for CMC = 17.85 μM. The magenta and green traces correspond to total A_4a_ and B_4b_ concentrations respectively. (E) Cluster size distributions at the last time point of CMC trajectories, that is 1000 ms for CMC = 16.7 μM, at 400ms for CMC = 17.85 μM and 18 μM. More detailed histograms without binning are shown in Supp. Fig. S6. (F) Total molecular concentrations converge to steady states as long as the solubility product (SP = 280 μM^2^) < Ksp and diverge with time when SP (= 330 μM^2^) > Ksp, irrespective of individual monomer concentrations.

Having established the Ksp with the FTC spatial simulations, we turned to the CMC approach, again using high values for k_create_ and k_decay_ (as in Fig. 2A). As we titrate up the CMCs (Fig. 7C), the system undergoes a fundamental change at 17.85 μM, corresponding to solubility product = 318.6 μM^2^, which is just above the Ksp we derived from the FTC calculations. Below that boundary (CMC = 16.7 μM), total concentrations converge to a steady state (Fig. 7C top panel and Movie S1); at (CMC = 17.85 μM) or above the boundary (CMC = 18 μM), they keep on going up with time (Fig. 7C 2^nd^ and 3^rd^ panels; Movie S2). When we look at the individual total concentration trajectories for CMC = 17.85 μM (Fig. 7D), much like our NFsim results, some trajectories fluctuate around a lower steady state (dilute phase) for the given timeframe while some trajectories shoot up (dense phase) after a variable lag. This stochastic behavior is also present for CMC = 18 μM (Supp. Fig. S7B), although the rate of growth is much faster in this case. Movie S2 dramatically illustrates how one trajectory reaches a metastable steady state, but ultimately explodes to fill up the available volume. All the trajectories fluctuate around a steady state for CMC = 16.7 μM (Supp. Fig. S7A; Movie S1), consistent with a single phase below the Ksp. Indeed, the molecular clusters, at the last time point of total concentration profiles, yield a bimodal distribution (Fig. 7E) above the Ksp while most molecules reside in small clusters below the phase boundary. So both FTC and CMC approaches are self consistent for our reference molecular pair: as the monomer SP remains below the Ksp threshold (318 μM^2^), the system exhibits only one phase; but above that threshold boundary, the molecules get partitioned into two different phases.

As for the NFSim model, the validity of the Ksp is preserved even when the CMCs are unequal. We illustrate this by choosing two solubility products, 280 μM^2^ and 330 μM^2^, respectively below and above the Ksp, and varying the individual CMCs (Fig. 7F). Both the cases with SP = 280 μM^2^ converge to a steady state, while SP = 330 μM^2^ combinations explode in both cases. It should be noted that as the total concentration explodes in these simulations, the volume can become filled with newly created molecules; this puts a brake on the total concentration in long duration simulations.

Next, we ask how molecular structures influence the phase boundary. We first consider a pair of tetravalent molecules with no linker sites (Fig. 8A, inset). Because the absence of linker sites reduces the degrees of freedom, we label this configuration a “low entropy system” or LES. With the FTC approach, we find that the Ksp for LES is much lower than the reference system of Fig. 7. However, the cluster size distributions show a dominant peak at the dimer (Supp. Fig. S8), both below and above the Ksp. To our surprise, when we clamp the monomer concentration at Ksp (CMC = 5.66 μM), the system does not show the characteristic diverging behavior with time (Fig. 8B). Even at CMC = 6 μM, total concentrations fluctuate around a steady state and show no sign of exploding over simulation times much longer than for the Reference system (compare Fig. 8B to Fig. 7C). Cluster size distributions at these two CMCs show a dominant dimer configuration compared to other molecular clusters (Fig. 8C). This huge “dimer trap” is a result of two biophysical factors – 1. low entropic cost upon forming the stoichiometrically matched dimer from less flexible monomers 2. perfect alignment of binding sites from two partner molecules with equal spacing between binding sites. This favorable dimeric state sucks up most of the monomers from solution and snaps them into a ladder like configuration where all the binding sites are saturated, preventing further recruitment of monomers to this dimer. Therefore, this “dimer trap” inhibits large cluster formation (no phase transition), but also reduces free monomer concentrations (Solubility products). The LES underscores the importance of structural flexibility and the role of the intrinsically disordered sequences in mediating phase transitions.

**Figure 8:**
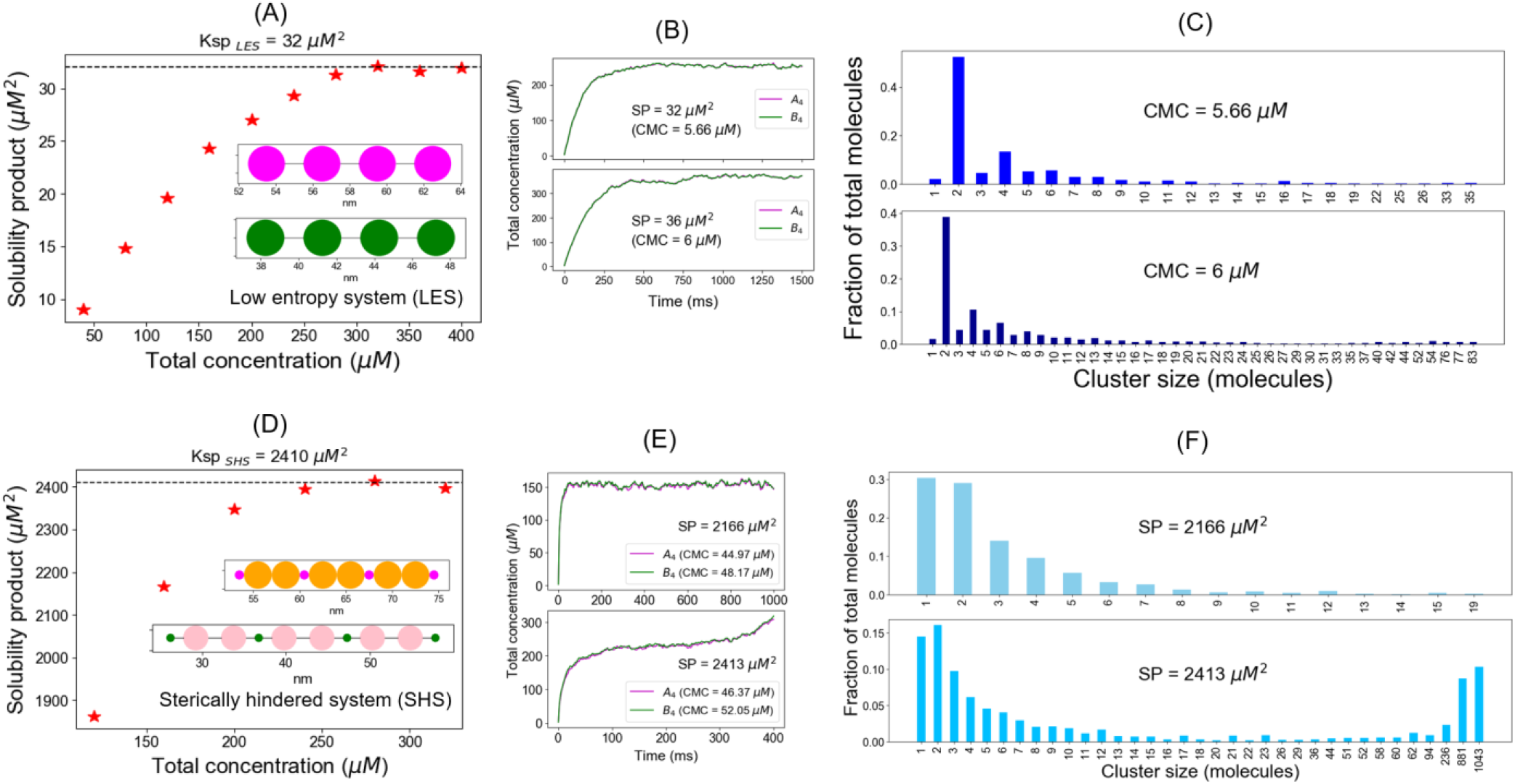
Effect of molecular structures on solubility product. (A) Solubility product profile of a pair of tetravalent molecules with no linker sites. Each of four magenta binding sites of A_4a_ can bind to one of the four green sites of B_4b_. Absence of linker sites reduce the degrees of rotational motion; hence we term this system as low entropy system or LES. Radius and diffusion constant of each site are 1 nm and 2 μm^2^/s respectively. For individual binding, dissociation constant, Kd = 350 μM. A total of 200 molecules (100 A_4a_ and 100 B_4b_) are placed in 3D boxes with varying volumes and monomer concentrations (averaged over 100 trajectories) are quantified. For LES, Ksp = 32 μM^2^, the horizontal dashed line. (B) Total molecular concentrations profile (averaged over 50 trajectories) at two different clamped monomer concentrations – one is just around Ksp (SP = 32 μM^2^) and another is well above Ksp (SP = 36 μM^2^). (C) Cluster size distributions (sampled from last time point) at two different CMCs. (D) Solubility product (SP) profile for a pair of molecules with four binding sites and unequal lengths. A_4a_ has four magenta binding sites (radius = 0.5 nm) and six orange linker sites (radius = 1.5 nm); B_4b_ has four green binding sites (radius = 0.5 nm) and six pink linker sites (radius = 1.5 nm). The bulky linker sites may interfere the binding between tiny binding sites; hence we term this system as sterically hindered system or SHS. Diffusion constant for each site is 2 μm^2^/s; dissociation constant for individual binding = 350 μM. Linear molecular length, A_4a_ = 21 nm and B_4b_ = 31.5 nm. Horizontal dashed line, Ksp = 2410 μM^2^. (E) Total molecular concentrations profile (averaged over 50 trajectories) at two different clamped monomer concentrations – one is below the Ksp (SP = 2166 μM^2^) and another is above Ksp (SP = 2413 μM^2^). (F) Cluster size distributions (sampled from last time point) at two CMCs.

To investigate the interplay between structural flexibility and inter binding site spacing, we create two pairs of rigid rod tetravalent binders. In one case, the spacing between binding sites are same (Supp. Fig. S9A) and in the other the spacing distance between binding sites is mismatched (Supp. Fig. S9B). Rigid binders with equal spacing yield a lower Ksp (Supp. Fig. S9C) and a stronger dimer trap, preventing large cluster formation (Supp. Fig. S9D). Unequal spacing between rigid binders breaks the dimer trap, pulling up the Ksp to a higher value and permitting the formation of large clusters (Supp. Fig. S9E). Thus a complex interplay between structural flexibility and geometrical arrangements of binding sites could play an important role in LLPS.

Next we seek to understand the role of steric interactions in phase transition. We create a pair of tetravalent binders where small binding sites are connected by bulkier linker sites – we term this configuration a sterically hindered system or SHS (Fig. 8D, inset). The lengths of these two flexible molecules are kept different to see whether that can lessen the dimer trap. The solubility product profile (Fig. 8D) yields a very high Ksp (~ 2410 μM^2^), suggesting inhibition of clustering by the unfavorable steric interference from the bulky linker sites. Also unequal molecular lengths result in different free concentrations for both the monomers (Supp. Fig. S10). However, clamping monomer concentrations above the Ksp does show diverging behavior with time (Fig. 8E) and a bimodal distribution of clusters (Figure 8F) despite having an abundance of monomer in the solution. The proportion of dimer is similar to the “Reference” system, the unequal spacing does not seem to diminish dimer formation, presumably because the molecules can fold into favorable conformations. The most important conclusion is that for this flexible system, steric hindrance around the binding sites increases the Ksp for LLPS, but does not prevent it.

## Discussion

It has been widely understood that for single component, self-interacting (homotypic) multivalent systems undergoing liquid-liquid phase separation (LLPS), the concentration of monomer in the dilute phase remains fixed above the phase transition, no matter how much of the monomer is added to the system; this feature of biomolecular condensates underlies buffering and noise reduction [4, 12]. Of course, the maintenance of a fixed concentration in a saturated solution of a solute in equilibrium with its solid phase is a feature of chemical systems that we all learned in high school. Recognizing this, we set out to see how far the analogy to solution chemistry could take us in considering heterotypic interactions between different multivalent binders. Somewhat more complex than the saturated solution of a single solute, is the precipitation of solid salts from saturated solutions of their ions. As we know from freshman chemistry (if not high school), ionic solutions of weakly soluble salts are governed by the solubility product constant, Ksp = [C^m+^]^n^[A^n−^]^m^, where m and n are the valencies, respectively, of the cation C^m+^ and anion A^n−^; importantly, n and m are the stoichiometries, respectively, of the cation and the anion in the solid phase because this stoichiometry matches the positive to the negative charges. We therefore explored theoretically, whether similar expressions could define the thresholds for LLPS in multicomponent (heterotypic) multivalent interaction systems.

Initially, we used a non-spatial stochastic network fee simulator, NFSim [16]. This allowed us to isolate only the effect of valency on the concentration dependence of clustering. The efficiency of this computational method also made it possible to screen many scenarios with a sufficient number of molecules and trajectories to assure statistically that we weren’t missing any interesting effects. We used 2 approaches to assess the threshold behavior. First, we titrated up the total concentration of pairs of multivalent binders. We found that there was a threshold above which the concentration of free monomers obeyed a Ksp expression – that is [A]^n^[B]^m^, where m and n are the ideal stoichiometries for an oligomer with all binding sites occupied. Below the Ksp threshold, the histogram of cluster size distributions tails off exponentially from monomers to small oligomers and its shape is independent of the number of molecules available (e.g Fig. 1B). However, for total concentrations above the Ksp threshold, the histogram of cluster size distributions becomes bimodal, with an increasing population of huge clusters as more molecules are added to the system (e.g. Fig. 1C); we consider this bimodal cluster size distribution to be a hallmark of phase separation. To further relate the behavior of these stochastic systems to the macroscopic phase transition, we used a second modeling approach where we clamped the free monomer concentrations (CMC) while allowing the clusters to grow. When monomer concentrations were clamped below the Ksp, the system reached a steady state identical to that of the corresponding closed fixed total concentration simulations, with identical cluster distributions containing a single decaying histogram of cluster sizes (Supp. Fig. S4). However, when the CMC was set to the Ksp or slightly above it, the plot of total concentration vs. time exploded following an initial lag (e.g. Fig. 2), indicating that as new monomers enter the system they were funneled into large clusters without attaining a steady state. Together, we feel that the behavior of these 2 different modeling approaches shows that the Ksp defines a threshold of monomer concentrations above which a phase separation occurs.

The generality of the Ksp as an indicator of threshold was then further tested using different scenarios. We extensively tested a situation where the total concentrations of each molecule in the binary tetravalent heterotypic system were unequal; while these simulations showed that the Ksp determined in the analysis of the equal concentration system was obeyed for a range of unequal concentrations (Fig. 3), there will be deviations if the range is extended too far (Supp. Fig. S2). This deviation is not surprising considering that the binding affinities are weak, leaving an excess of binding sites empty within a cluster, when the free concentration of one binding partner is too highly depleted. We tested cases where the valencies were not identical for a binary (Fig. 5) and even a ternary system of mixed valency interactors (Fig. 6). In these systems, a more complex Ksp formulation was required to account for the appropriate ideal stoichiometry of the cluster. In both of these cases, the Ksp was successful in defining a threshold for phase separation.

We turned to coarse grained spatial simulations of clustering to determine if the Ksp might still be generally applicable when the shape and flexibility of multivalent molecules is explicitly considered. We used SpringSaLaD [18] software; it models molecules as a series of linked spheres to represent domains within macromolecules. We started with a heterotypic pair of tetravalent binders (Fig.7), similar to the non-spatial model used in Figs. 1 and 2. The SpringSaLaD structures included 4 binding spheres, each separated by pair of linker spheres; the distances between sites were identical within each binding partner. The results (Fig. 7) show that this model displays the same behavior noted for the non-spatial model. The product of concentrations of monomer becomes independent of the total concentration above a threshold (Fig. 7B). When monomer concentrations are clamped at or above this Ksp threshold, the total concentration explodes after a lag time (Figs 7C and F and Movie S2) and the histogram of cluster sizes becomes bimodal (Fig. 7E). Thus, the Ksp concept seems to apply to at least the structure shown in Fig. 7A.

The SpringSaLaD models allow us to explore which molecular structural features may be required for the Ksp to be able to control phase separation; we could also ask how the structures of the interacting multivalent molecules might influence the value of the Ksp, which is a measure of the tendency of a system toward phase separation. We used the system in Fig. 7A as a reference baseline, varying one structural feature at a time. As an example, but not surprisingly, increasing the size of the linker spheres while decreasing the size of the binding site spheres increases the Ksp rather drastically (Fig. 8D). So steric effects can alter the tendency of these systems to cluster. But, importantly, the Ksp still marks the threshold for phase separation (Figs. 8E and F).

In contrast, removal of linker sites does break the ability of Ksp to define a threshold for phase separation (Figs. 8A, B, C). There is an *apparent* Ksp that represents a limit of monomer concentration above a threshold total concentration (8A); however, total concentrations above this threshold do not produce a bimodal cluster histogram (8C), nor, in the CMC simulations, a total concentration that explodes with time (8B). While the 4 spheres in the molecules of Fig. 8A are free to adopt different angles and dihedral angles with a fixed bond length between them, the molecules have fewer degrees of freedom than the structures in Fig. 7A, with their additional linker spheres. So, the system in Fig. 8A has relatively lower entropy than the system of Fig. 7A. Therefore, the formation of a fully bound dimer between 2 heterotypic monomers would lose less entropy for the structures in 8A compared to 7A. Another equivalent way of thinking about this is that the molecules in 8A can more easily align their binding sites to match the positions of the complementary binding sites to form the dimer. We called this tendency to form dimers, evident in the histograms of Fig. 8C, a “dimer trap”, because it funnels most monomers into the dimer state and prevents the formation of larger clusters, even when the monomer concentrations are clamped considerably above the Ksp. The tendency to form dimers also exists for the reference structures in Fig. 7A, as shown in Supp. Fig. S6. The 7A structures also have perfect spatial matching of binding sites, but the additional entropy imparted by the flexible linker domains assures that the dimer is not so dominant as to completely overpower the tendency to form large clusters.

This perfect matching of binding sites has been previously noted in rigid body simulations as an impediment to the growth of large clusters [20, 21] and we were able to reproduce this with the rigid linear structures in Supp. Fig. S9. When the spacing between binding sites matches, these rigid systems fall into a dimer trap and don’t produce a bimodal cluster histogram (panel 9D) until well beyond threshold concentrations for an apparent Ksp (panel 9C); if the spacing is mismatched, the rigid system does form large clusters at lower concentrations (panel 9E), but the Ksp is poorly defined (panel 9C). Thus, from these computational experiments it appears that heterotypic binders with limited conformational freedom may not be governed by the Ksp formulation and may not be likely to undergo LLPS. This is consistent with the established experimental and theoretical result that robust LLPS requires that the binding sites in multivalent molecules be interspersed with disordered protein regions [22–24]. On the other hand, spatial matching of binding sites and limited flexibility may be the hallmarks for assembly of stoichiometric molecular machines as opposed to pleomorphic condensates [14].

Manipulation of computational models thus allowed us to systematically determine how well the Ksp concept, familiar from ionic solution chemistry, might apply to the very different situation of weak interactions between multivalent macromolecules. We found that for most scenarios, if not all, the Ksp does define a threshold for the unbounded growth of large clusters and a quantitative metric for the tendency of a system to phase separate. Additionally, for those cases where there are deviations from the stereotypical behavior, insights may be gleaned into the underlying molecular factors inhibiting cluster growth. But it is important to appreciate that experimental systems may introduce additional levels of complexity, such as nonspecific low affinity binding interactions and long-range electrostatic forces. For real biomolecular condensates, the valency and stoichiometry of the components may be unknown, and they may have multiple binding interaction of differing affinities. However, it should be possible to measure Ksp experimentally using fluorescently labelled binding partners and determining their concentrations in the low-density phase. Such measurements could actually provide some indication of the effective valency of the interactions in complex multivalent molecular ensembles.

## Supporting information

unbounded growth of molecular clusters above ksp

Molecular clusters approach to a steady state below Ksp

## Acknowledgments

We are sincerely thankful to Dr. Boris Slepchenko for the thoughtful discussions on clamped monomer concentration approach. We gratefully acknowledge support of this work by NIH grants R01 GM132859 and R24 GM137787 from the National Institute for General Medical Sciences.

## Reference

1. Banani, S.F., et al., Biomolecular condensates: organizers of cellular biochemistry. Nature Reviews Molecular Cell Biology, 2017. 18: p. 285.

2. Hyman, A.A., C.A. Weber, and F. Jülicher, Liquid-Liquid Phase Separation in Biology. Annual Review of Cell and Developmental Biology, 2014. 30(1): p. 39–58.

3. Shin, Y. and C.P. Brangwynne, Liquid phase condensation in cell physiology and disease. Science, 2017. 357(6357).

4. Holehouse, A.S. and R.V. Pappu, Functional Implications of Intracellular Phase Transitions. Biochemistry, 2018. 57(17): p. 2415–2423.

5. Su, X., et al., Phase separation of signaling molecules promotes T cell receptor signal transduction. Science (New York, N.Y.), 2016. 352(6285): p. 595–599.

6. Shin, Y. and C.P. Brangwynne, Liquid phase condensation in cell physiology and disease. Science, 2017. 357(6357): p. eaaf4382.

7. Alberti, S., A. Gladfelter, and T. Mittag, Considerations and Challenges in Studying Liquid-Liquid Phase Separation and Biomolecular Condensates. Cell, 2019. 176(3): p. 419–434.

8. Mathieu, C., R.V. Pappu, and J.P. Taylor, Beyond aggregation: Pathological phase transitions in neurodegenerative disease. Science, 2020. 370(6512): p. 56.

9. Li, P., et al., Phase transitions in the assembly of multivalent signalling proteins. Nature, 2012. 483: p. 336.

10. Shin, Y., et al., Spatiotemporal Control of Intracellular Phase Transitions Using Light-Activated optoDroplets. Cell, 2017. 168(1): p. 159–171.e14.

11. Wang, J., et al., A Molecular Grammar Governing the Driving Forces for Phase Separation of Prion-like RNA Binding Proteins. Cell, 2018. 174(3): p. 688–699.e16.

12. Klosin, A., et al., Phase separation provides a mechanism to reduce noise in cells. Science, 2020. 367(6476): p. 464.

13. Riback, J.A., et al., Composition-dependent thermodynamics of intracellular phase separation. Nature, 2020. 581(7807): p. 209–214.

14. Mayer, B.J., M.L. Blinov, and L.M. Loew, Molecular machines or pleiomorphic ensembles: signaling complexes revisited. J Biol, 2009. 8(9): p. 81.1–81.8.

15. Falkenberg, Cibele V., Michael L. Blinov, and Leslie M. Loew, Pleomorphic Ensembles: Formation of Large Clusters Composed of Weakly Interacting Multivalent Molecules. Biophys J, 2013. 105(11): p. 2451–2460.

16. Sneddon, M.W., J.R. Faeder, and T. Emonet, Efficient modeling, simulation and coarse-graining of biological complexity with NFsim. Nature Methods, 2011. 8(2): p. 177–183.

17. Chattaraj, A., M. Youngstrom, and L.M. Loew, The Interplay of Structural and Cellular Biophysics Controls Clustering of Multivalent Molecules. Biophys J, 2019. 116(3): p. 560–572.

18. Michalski, P.J. and L.M. Loew, SpringSaLaD: A Spatial, Particle-Based Biochemical Simulation Platform with Excluded Volume. Biophysical journal, 2016. 110(3): p. 523–529.

19. Posey, A.E., A.S. Holehouse, and R.V. Pappu, Chapter One - Phase Separation of Intrinsically Disordered Proteins, in Methods in Enzymology, E. Rhoades, Editor. 2018, Academic Press. p. 1–30.

20. Freeman Rosenzweig, E.S., et al., The Eukaryotic CO2-Concentrating Organelle Is Liquid-like and Exhibits Dynamic Reorganization. Cell, 2017. 171(1): p. 148–162.e19.

21. Xu, B., et al., Rigidity enhances a magic-number effect in polymer phase separation. Nature Communications, 2020. 11(1): p. 1561.

22. Choi, J.-M., A.S. Holehouse, and R.V. Pappu, Physical Principles Underlying the Complex Biology of Intracellular Phase Transitions. Annual Review of Biophysics, 2020. 49(1): p. 107–133.

23. Harmon, T.S., et al., Intrinsically disordered linkers determine the interplay between phase separation and gelation in multivalent proteins. eLife, 2017. 6: p. e30294.

24. Pak, Chi W., et al., Sequence Determinants of Intracellular Phase Separation by Complex Coacervation of a Disordered Protein. Molecular Cell, 2016. 63(1): p. 72–85.

