## Supplementary figures for "Solubility product constant directs the formation of biomolecular condensates"

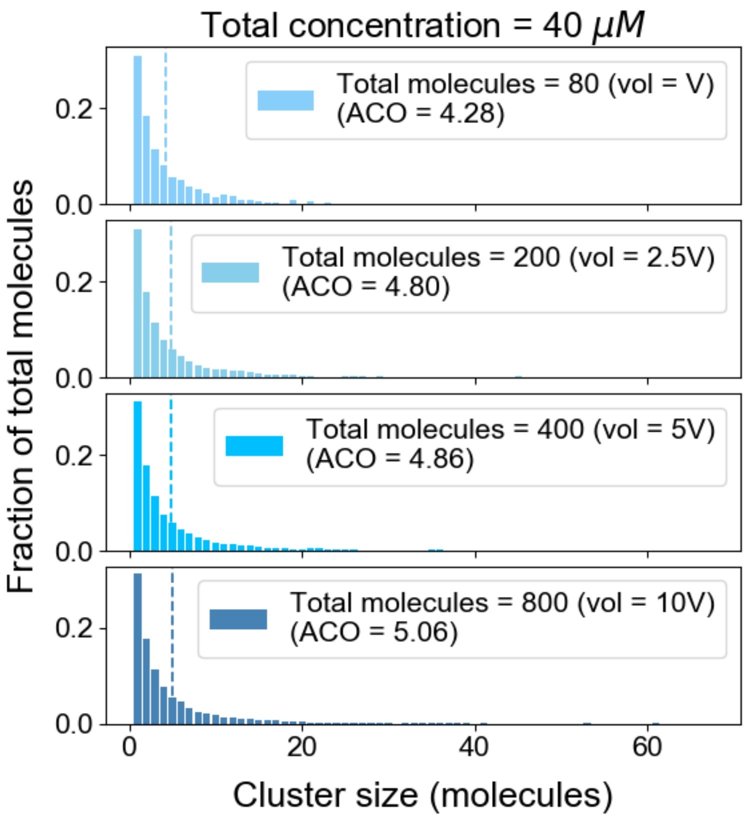


Figure S1: System shows similar cluster size distributions with varying volume and monomer counts as long as the total concentration (40 µM in this case) is below the threshold concentration (~ 60 µM for our $A_4$ – $B_4$ system). We term the mean of the distribution as average cluster occupancy (ACO) which is shown with dotted line in each case.

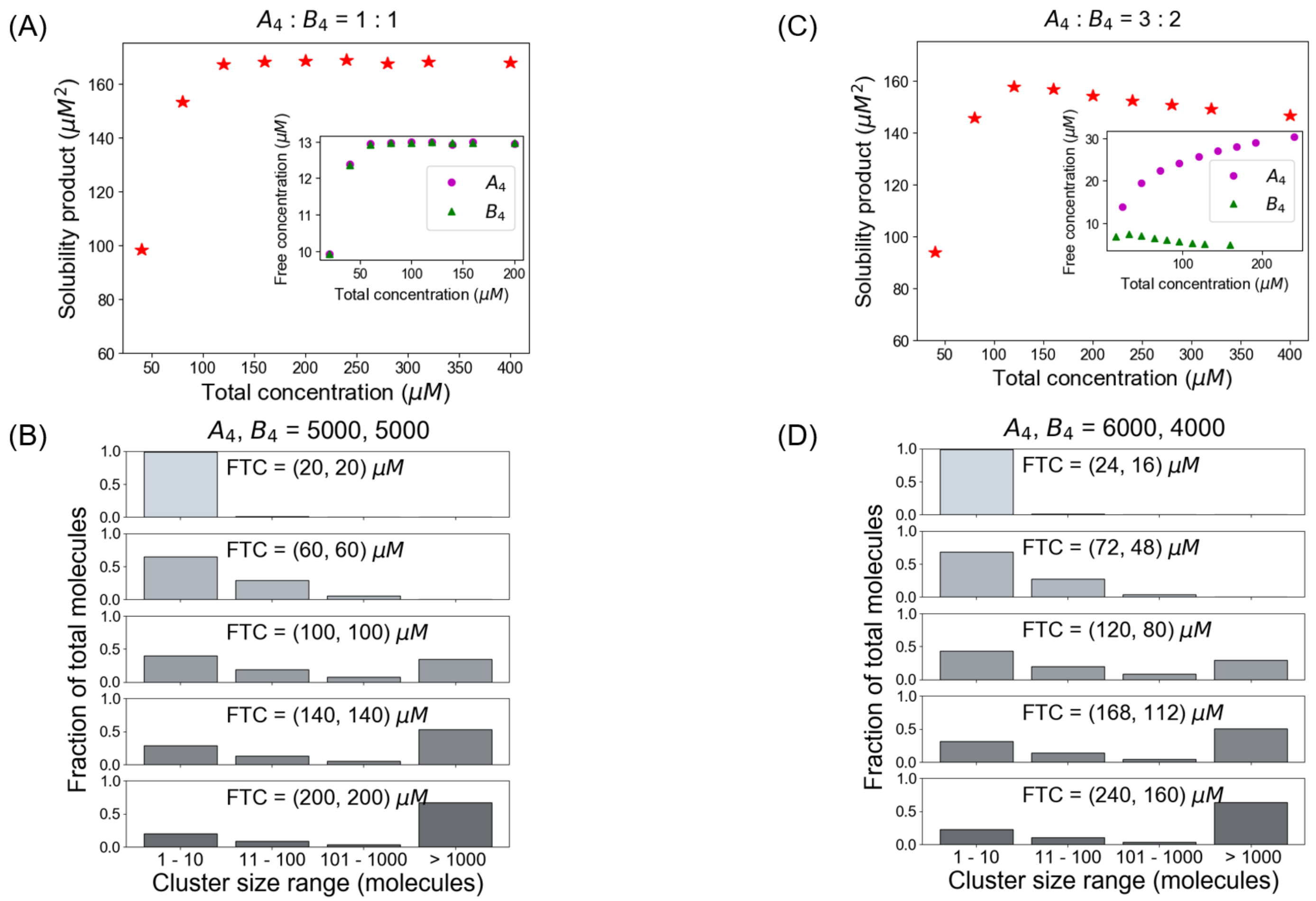


Figure S2: System deviates from a fixed Ksp when initial conditions are not stoichiometrically matched. (A, B) Solubility product profile and cluster size distributions when we use a pair of tetravalent molecules ($A_4$ and $B_4$) in equal stoichiometric amount. In all these cases, 5000 $A_4$ and 5000 $B_4$ molecules are kept fixed and we vary the Kd to change the concentrations. (C, D) Solubility product profile and cluster size distributions with unequal stoichiometric ratio. In all these simulations, we maintain a fixed number of 6000 $A_4$ and 4000 $B_4$ molecules and change the Kd to alter the concentrations.

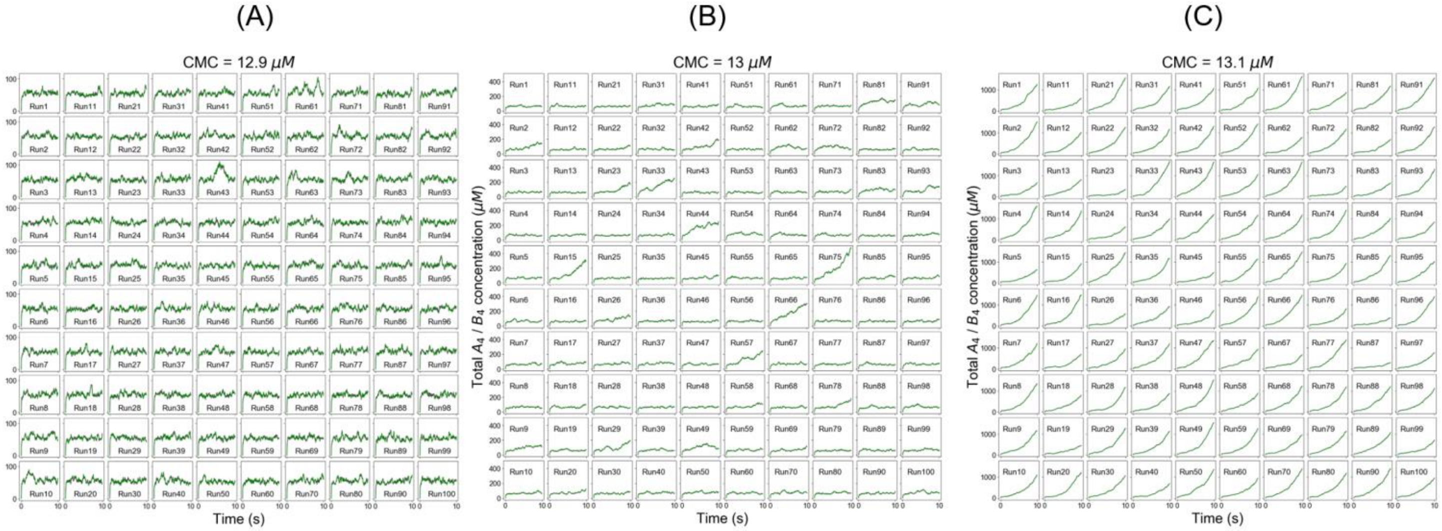


Figure S3: Systems behaves more stochastically at the phase boundary. Individual trajectories of total molecular concentrations ($A_4$ in magenta and $B_4$ in green) at (A) CMC = 12.9 µM (B) CMC = 13 µM (C) CMC = 13.1 µM. All the trajectories show similar trends for CMC = 12.9 µM (fluctuation around a steady state) or CMC = 13.1 µM (diverging growth with time). However, for CMC = 13 µM, where solubility product = 13*13 $\mu M^2$ = 169 $\mu M^2$, system behaves quite randomly across different trajectories; some of them seem to be fluctuating around a steady state while some of the trajectories show the diverging behavior.

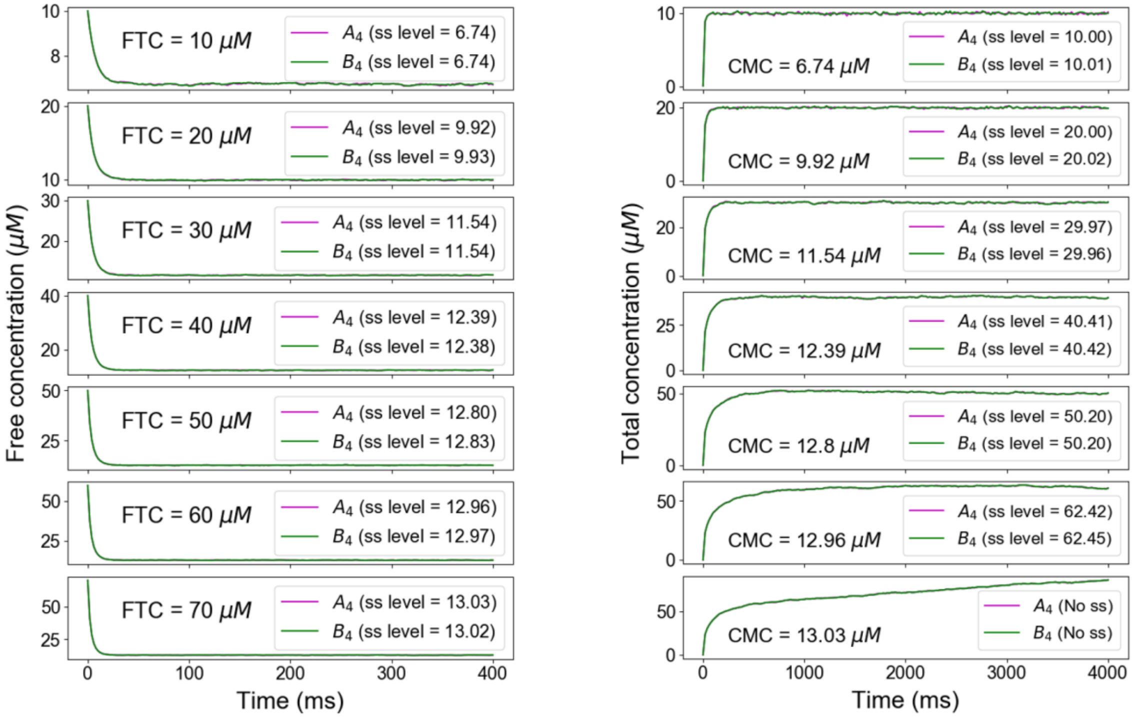


Figure S4: Summary of FTC and CMC method predictions. Below the phase boundary, if we clamp the free concentrations from the FTC approach, we recover the identical total concentrations at steady state in CMC approach. Above the concentration threshold (example of FTC = 70 μM), clamping the free concentrations from FTC method no longer yields the same steady state level (CMC = 13.01 μM).

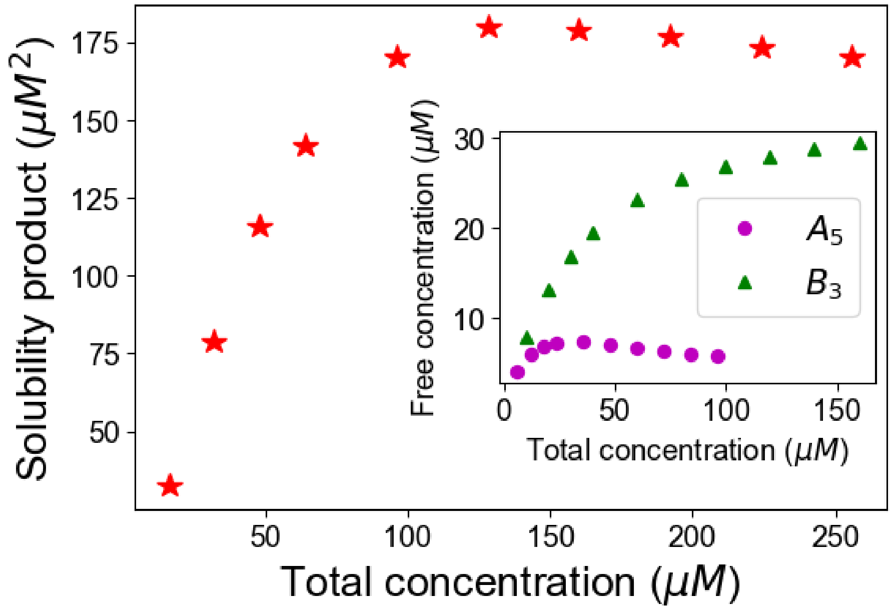


Figure S5: Simple product of free molecular concentrations does not work for mixed valent binary systems. Here, solubility product = Free $A_5$ * Free $B_3$

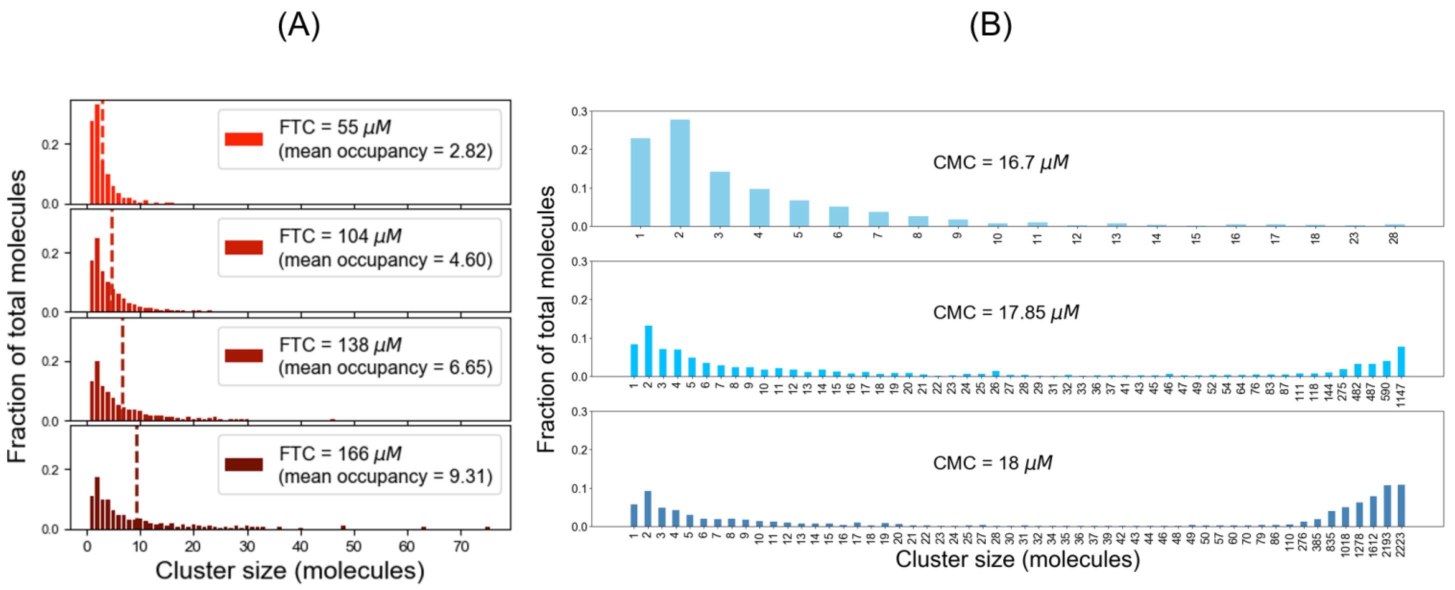


Figure S6: For reference system, (A) cluster size distributions at steady state (sampled from last time point) for different fixed total concentrations; (B) Cluster size distribution at last time point for different clamped monomer concentrations.

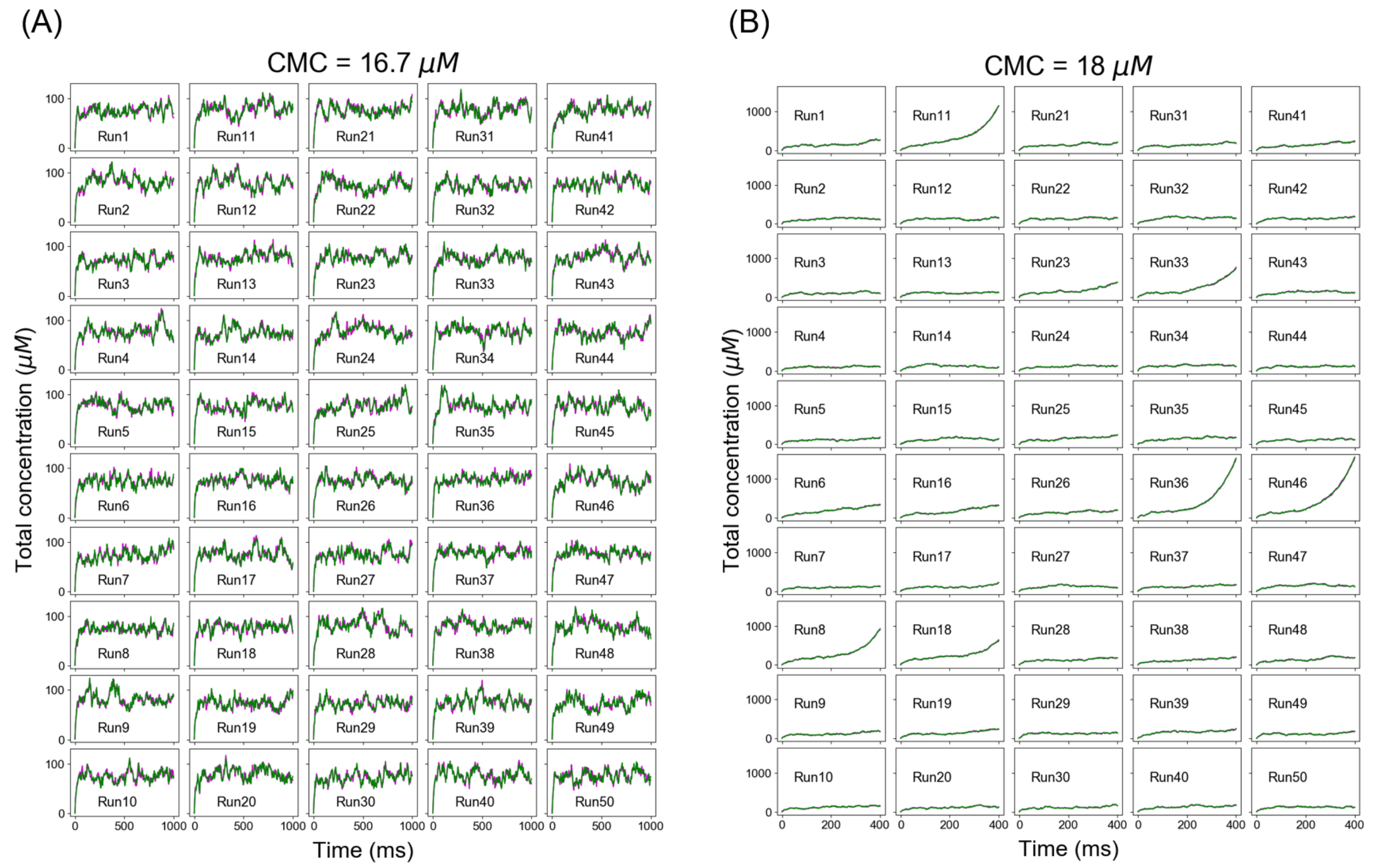


Figure S7: For reference system, individual total concentration trajectories at (A) CMC = 16.7 μM (below Ksp), and (B) CMC = 18 μM (above Ksp). Magenta and green traces correspond to total $A_4$ and total $B_4$ concentration respectively.

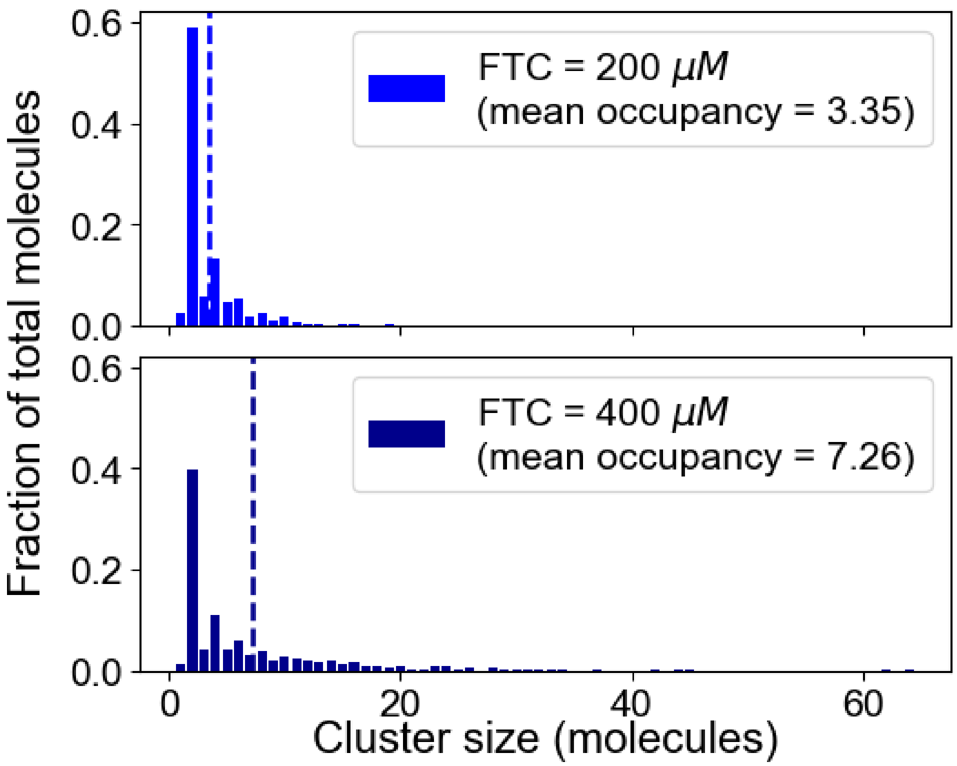


Figure S8: Cluster size distributions at two different fixed total concentrations for low entropy system.

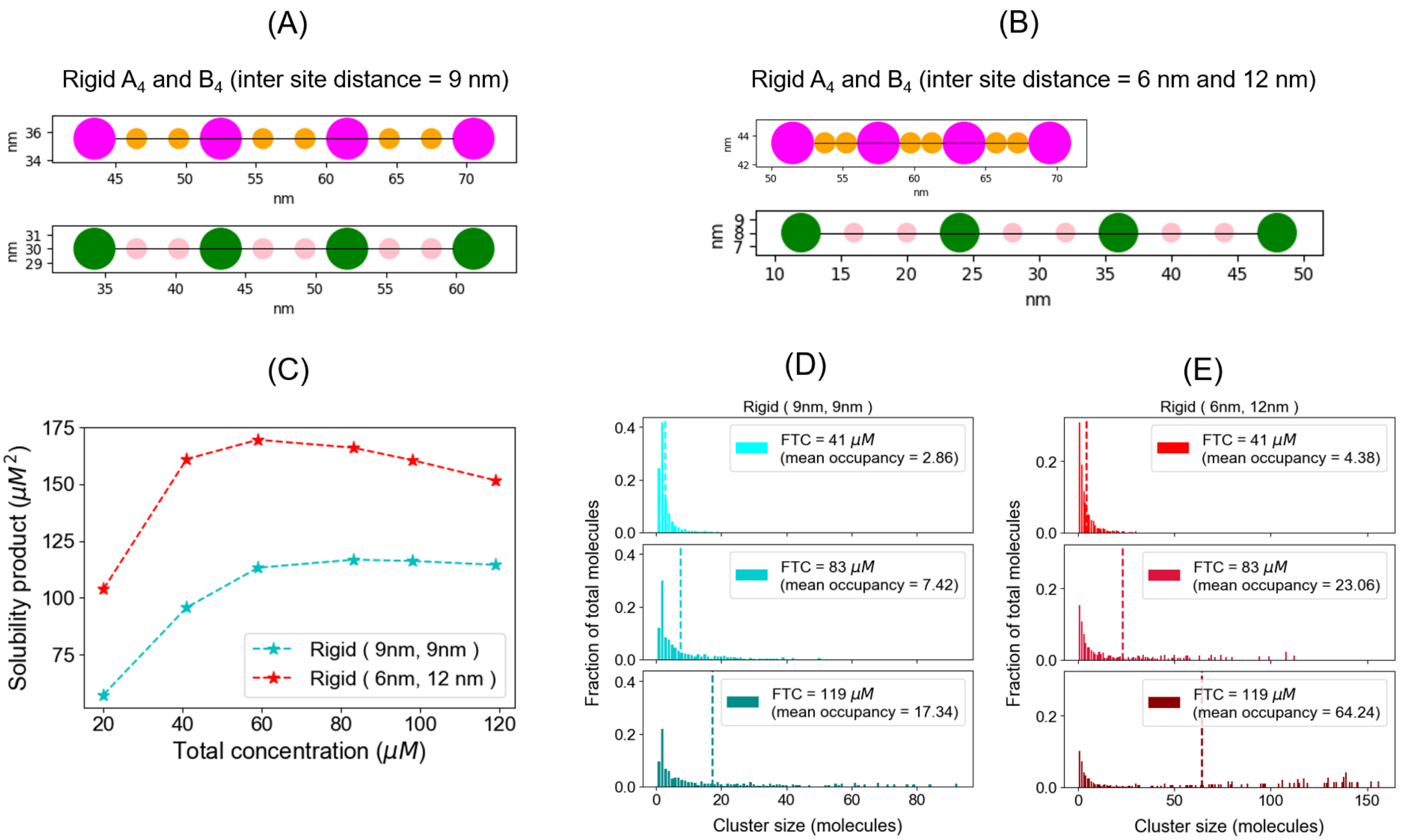


Figure S9: Interplay of molecular rigidity and inter binding site distance regulates dimer trap. (A) A tetravalent molecular pair (rigid rod like) with equal spacing between binding sites where distance between two binding sites (magenta or green) is 9 nm. Radius, magenta and green sites = 1.5 nm; orange and pink sites = 0.75 nm. Diffusion constants for all the sites are 2 $\mu m^2/s$. (B) A tetravalent rigid molecular pair with unequal spacing between binding sites, where distances between two magenta and green sites are 6 nm and 12 nm respectively. Radius, magenta and green sites = 1.5 nm; orange and pink sites = 0.75 nm. Diffusion constants for all the sites are 2 $\mu m^2/s$. (C) Solubility product profiles for configuration A (cyan) and B (red). Total 200 molecules (100 $A_4$ and 100 $B_4$) are placed in 3D boxes with varying volumes and free molecular concentrations are quantified. Dissociation constant for individual binding = 350 μM for both the systems. (D, E) Cluster size distributions at three different fixed total concentrations (sampled from last time point) for configuration A and B.

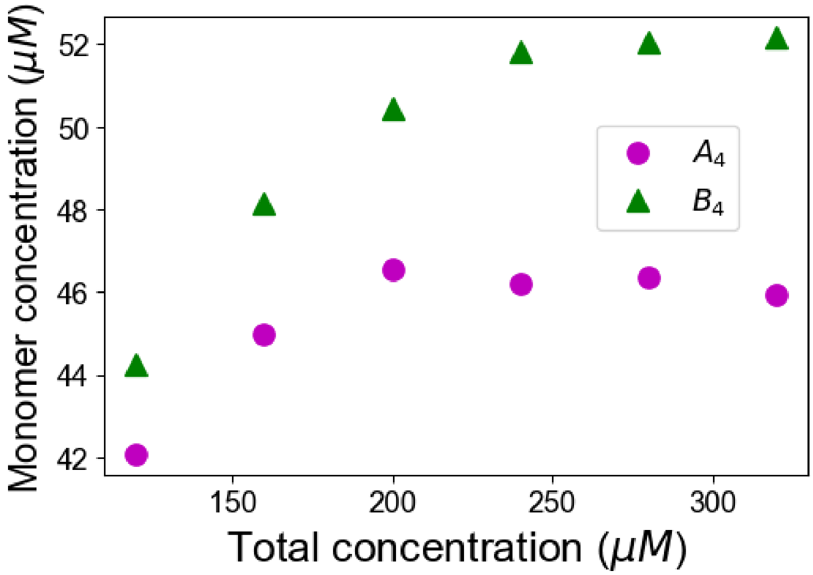


Figure S10: For sterically hindered system, free molecular concentrations as a function of total concentrations.
